# INfrastructure for a PHAge REference Database: Identification of large-scale biases in the current collection of phage genomes

**DOI:** 10.1101/2021.05.01.442102

**Authors:** Ryan Cook, Nathan Brown, Tamsin Redgwell, Branko Rihtman, Megan Barnes, Martha Clokie, Dov J. Stekel, Jon Hobman, Michael A. Jones, Andrew Millard

## Abstract

**Background:** With advances in sequencing technology and decreasing costs, the number of bacteriophage genomes that have been sequenced has increased markedly in the last decade.

**Materials and Methods:** We developed an automated retrieval and analysis system for bacteriophage genomes, INPHARED (https://github.com/RyanCook94/inphared), that provides data in a consistent format.

**Results:** As of January 2021, 14,244 complete phage genomes have been sequenced. The data set is dominated by phages that infect a small number of bacterial genera, with 75% of phages isolated only on 30 bacterial genera. There is further bias with significantly more lytic phage genomes than temperate within the database, resulting in ~54% of temperate phage genomes originating from just three host genera. Within phage genomes, putative antibiotic resistance genes were found in higher frequencies in temperate phages than lytic phages.

**Conclusion:** We provide a mechanism to reproducibly extract complete phage genomes and highlight some of the biases within this data, that underpins our current understanding of phage genomes.

## Introduction

Bacteriophages (hereafter phages), are viruses that specifically infect bacteria and are thought to be the most abundant biological entities in the biosphere (1). In the oceans they are important in diverting the flow of carbon into dissolved and particulate organic matter via the lysis of their hosts (1), or directly halting the fixation of CO_2_ carried out by their cyanobacterial hosts (2). In the human microbiome, it is becoming increasingly clear that phages play a role in a range of different diseases. Many recent studies have shown disease-specific alterations to the gut virome community in both gastrointestinal and systemic conditions, including irritable bowel disease (3), AIDs (4), malnutrition (5), and diabetes (6).

Phages alter the physiology of their hosts such that their bacterial hosts display increased virulence, a notable example being phage CTX into the genome of *Vibrio cholerae*, resulting in cholera (7). However, there are many cases where the expression of phage-encoded toxins cause an otherwise harmless commensal bacterium to convert into a pathogen, including multi-drug resistant ST11 strains of *Pseudomonas aeruginosa* (8, 9), and the Shiga-toxin encoding *Escherichia coli* (10). As well as increasing the virulence of the host bacteria, phages can also utilise parts of their genomes known as auxiliary metabolic genes (AMGs), homologues of host metabolic genes, to modulate their hosts metabolism (11).

Our understanding of how phages alter host metabolism has increased in conjunction with the number of phage genomes that have been sequenced, following sequencing of the first phage genome in 1977 (12). Since then, the number of phages that are isolated and the relative ease of high-throughput sequencing has led to a rapid increase in the number of sequenced bacteriophage genomes (13). The relatively simple nature of phage genomes means that the vast majority of isolated phage genomes can be completely assembled using short-read next generation sequencing (14). The greater number of phage genomes available results in common analyses, including comparative genomic analyses (15, 16), taxonomic classification of phages (17–20), forming the basis of software to predict new phages (21–26), and as is often the first step in analysis of viromes, the comparison of sequences to a known database.

To do all of the above requires a comprehensive set of complete phage genomes from cultured isolates that can be used to build databases for further analyses. It also raises the question of how many complete phage genomes are currently available. While this should be relatively trivial question to answer, it is not very simple to do so, as there is currently no such database of all complete phage genomes. Therefore, the aim of this work was to provide a reproducible and automated way to extract complete phage genomes from GenBank and identify general properties within the data and limitations.

## Materials and Methods

Bacteriophage genomes were download using the “PHG” identifier along with minimum and maximum length cut-offs. Genomes were then filtered based on several parameters to identify complete and near complete phage genomes. This includes initial searching for the term “Complete” & “Genome” in the phage description, followed by “Complete” & (“Genome” or “Sequence”) & a genome length of greater than 10 kb. The list of genomes was then manually curated to identify obviously incomplete phage genomes, with the process on going. The accessions of these are then excluded in future iterations by the use of an exclusion list, which can be added to by the community via GitHub. Whilst this process is not perfect, we thank numerous people that have identified genomes within this list that are obviously incomplete. The initial search term for downloading genomes was: *esearch - db nucleotide -query “gbdiv_PHG[prop]” | efilter -query “1417:800000 [SLEN]” | efetch - format gb > $phage_db.gb*. An exclusion list of phage genomes that are automatically called “complete”, yet when manually checked are not is continually being updated.

After filtering, genes are called using Prokka with the ‒noanno flag, with a small number of phages using ‒gcode 15 (27, 28). Gene calling was repeated to provide consistency across all genomes, which is essential for comparative genomics. A database is provided so that this process does not continually have to be rerun and only new genomes are added. The original GenBank files are used to gather useful metadata including taxa and bacterial host, and the Prokka output files are used to gather data relating to genomic features. The gathered data are summarised in a tab-delimited file that includes the following: accession number, description of the phage genome, GenBank classification, genome length (bp), molGC (%), modification date, number of CDS, proportion of CDS on positive sense strand (%), proportion of CDS on negative sense strand (%), coding capacity (%), number of tRNAs, bacterial host, viral genus, viral sub-family, viral family, and the lowest viral taxa available (from genus, sub-family and family). Coding capacity was calculated by comparing the genome length to the sum length of all coding features within the Prokka output, and tRNAs were identified by the use of tRNA tag. Other outputs include a fasta file of all phage genomes, a MASH index for rapid comparison of new sequences, vConTACT2 input files, and various annotation files for IToL and vConTACT2. The vConTACT2 input files produced from the script were processed using vConTACT2 v0.9.13 with --rel-mode Diamond --db ‘None’ --pcs-mode MCL --vcs-mode ClusterONE --min-size 1 and the resultant network was visualised using Cytoscape v3.8.0 (29, 30).

To identify genes indicative of a temperate lifestyle within genomes, we used a set of PFAM HMMs as described previously (31). If a genome encoded one of these genes, it was assumed to be temperate. Antimicrobial resistance genes (ARGs) and virulence factors were identified using Abricate with the resfinder and VFDB databases using 95% identity and 75% coverage cut-offs (32–34).

The phylogeny of “jumbo-phages” was constructed from the amino acid sequence of the TerL protein, extracted from 313/314 of the “jumbo-phage” genomes. Sequences were queried against a database of proteins from non-“jumbo-phages” using Blastp and the top five hits were extracted (35) with redundant sequences being removed. Sequences were aligned with MAFFT, with a phylogenetic tree being produced using IQ-Tree with “-m WAG - bb 1000” which was visualised using IToL (36–38). Additional information was overlaid using IToL templates that are generated via INPHARED.

Rarefaction analysis was carried out for phage genomes from the top ten most common hosts (70% ID over 95% length) and species (95% ID over 95% length) using ClusterGenomes v5.1 (39). An additional set of these genomes pooled together was included. Rarefaction curves and species richness estimates were produced using Vegan in R (40, 41).

All data from January 2021 is available at Figshare https://doi.org/10.25392/leicester.data.14242085 and the script used for downloading and analysing genomes is available on GitHub https://github.com/RyanCook94/.

## Results

The output of the INPHARED script provides as set of complete phage genomes, whereby genes have been called in a consistent manner that allows comparative genomics and phylogenetic analysis. In addition, it provides a MASH database to allow rapid comparison of new phage genomes against to identify close relatives. Along with formatted databases for input into vConTACT2 to allow identification of more distant relatives. The host data (Genus) for each phage is extracted along with summary information for each genome, which is reformatted to allow overlay onto trees in IToL (See Supplementary Figure 1 for full details).

For this study, we used a lenient definition of “complete” for the identification of complete phage genomes. Strictly speaking a complete phage genome would include the terminal ends of the phage genome. As many phages are sequenced using a transposon based library preparation (16, 42), the genome can never be complete as the terminal bases can never be sequenced, unless it is circularly permuted (14). For phage genomes with long terminal repeats, if the length of the repeat is larger than the library insert size, these cannot be resolved. As this information is not included in every GenBank file, automated retrieval is not possible.

We next set about identifying how many phage genomes have been sequenced to date. The extraction of genomes from the nucleotide database of GenBank results in 18,134 genomes. Of these, 3,890 phage genomes are REFSEQ entries which are derived from primary submissions, resulting in 14,244 putative complete phage genomes. Current recommendations are that phages are uniquely named (43), if this assumption is true then number of unique phage genomes is 12,127 if phages with the same name are truly identical. However, there are multiple examples of phages with the same name. In some cases this is the same phage being re-sequenced due to experimental evolution studies such as *E. coli* phiX (44). In other instances, phages with the same name are not genetically identical. Thus, using a phage name as a means to identify different phages is not a suitable method for determining the number of unique phage genomes. As an alternative, de-duplication of genomes at 100%, 97% and 95% identity results in 13,830, 12,845 and 12,770 genomes respectively.

Having established a dataset of “complete” phage genomes, we then analysed this data to look at general phage genomic properties. First, we looked at the increase in the number of phage genomes that are sequenced over time. Whilst the number of phage genomes has rapidly increased over the last 20 years, the rate of increase has slowed in the last decade (Figure 1), with the number of phage genomes doubling every 2-3 years.

**Figure 1.**
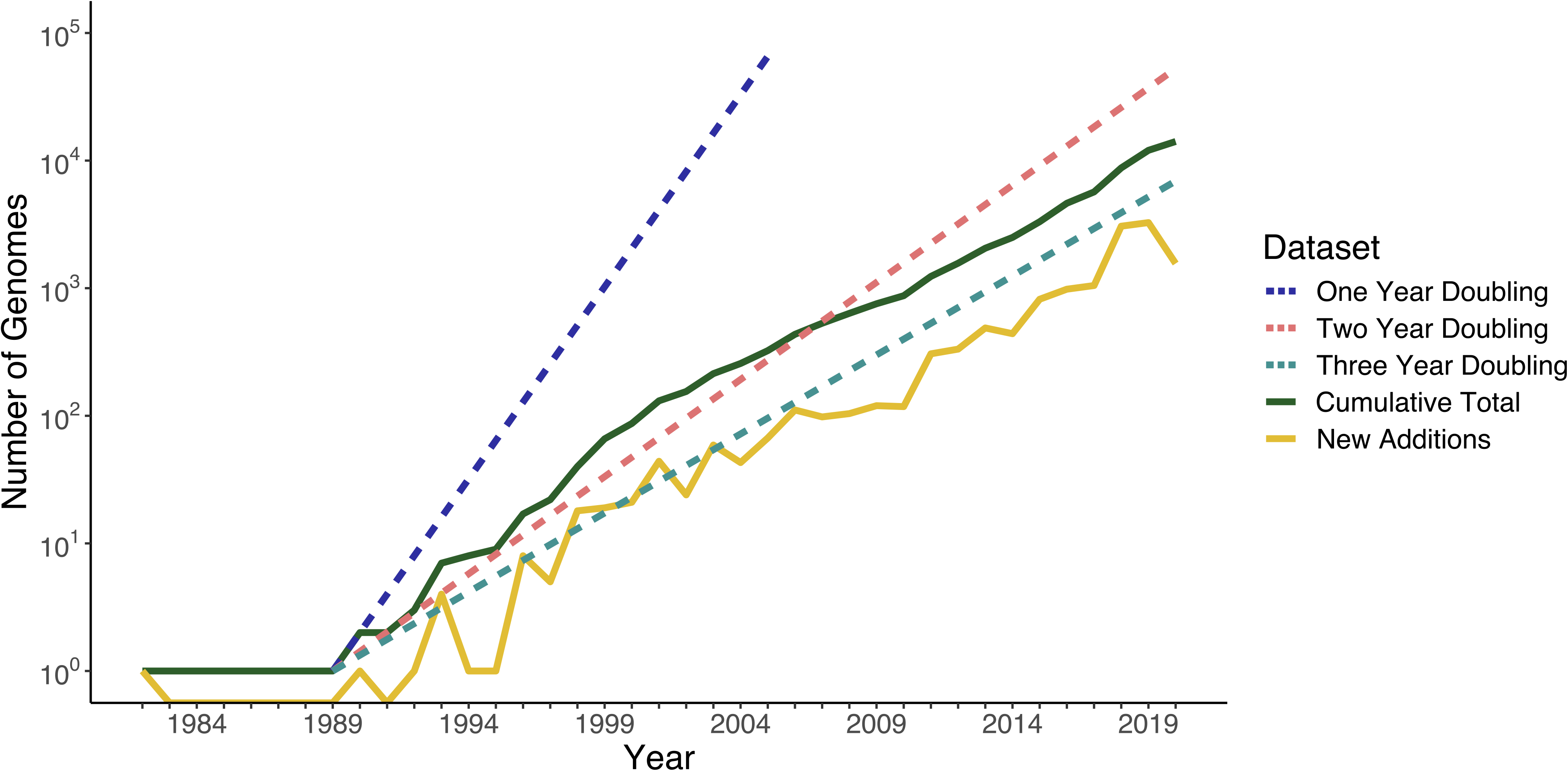
Number of complete phage genomes in GenBank over time. Dates were estimated based on date of submission (* for 235 genomes, the date of update was used as no submission date was available). The reference lines showing doubling rates (dashed) begin in 1989, as this is when the number of phage genomes increased beyond the first submission in 1982.

### Bacteriophage Hosts and Predicted Gene Function

Utilising the database of complete genomes, we extracted the hosts and predicted number of hypothetical proteins for each phage. Across all phages, the majority of genes which encode proteins with unknown function (hypothetical) was mean 56% (+/− 20), supporting the truism that the majority of genes encode proteins within unknown function.

The host of 12,403 phages were extracted with the remainder unknown as the host was not clear from the information contained within the GenBank file alone. The genomes of phages infecting 234 hosts have been sequenced. However, there is a clear bias in the isolation of phages against the same host (Figure 2a). Phages that infect *Mycobacterium* spp. are the most commonly deposited genomes (~13%), largely due to the pioneering work of the SEA-PHAGES program (45), followed by *Escherichia* spp., *Streptococcus* spp., and *Pseudomonas* spp. (Figure 2a). Phages isolated on just 30 different bacterial genera accounts for ~75% of all phage genomes in the database (Supplementary Table 1). For genomes isolated against the top ten hosts, we used rarefaction analysis to gain an understanding of the diversity of phage genomes isolated to date and determine redundancy of phages isolated on a particular host. Using a cut-off of 95% identity to define a species, it was clearly observed the number of phage species continues to increase with the number of genomes sequenced, a pattern also observed at the level of genus (70% identity) (Figure 3). Using the current data, it was possible to estimate how many different species of phage might infect these different hosts (Supplementary Table 4). For *Mycobacterium*, for which most phages have isolated on, there are 695 observed species with an estimated 2132-2282 total species. Thus, demonstrating even for hosts where thousands of phages have been isolated, we are only just scratching the surface of the diversity of total phage diversity. We are also likely under estimating the total number of different phage species. In the case of phages infecting *Mycobacterium*, the majority of these have been isolated on only a single strain as part of the SEA-PHAGES program. Increasing the diversity of the host *Mycobacterium*, is likely to lead to higher estimates.

**Figure 2.**
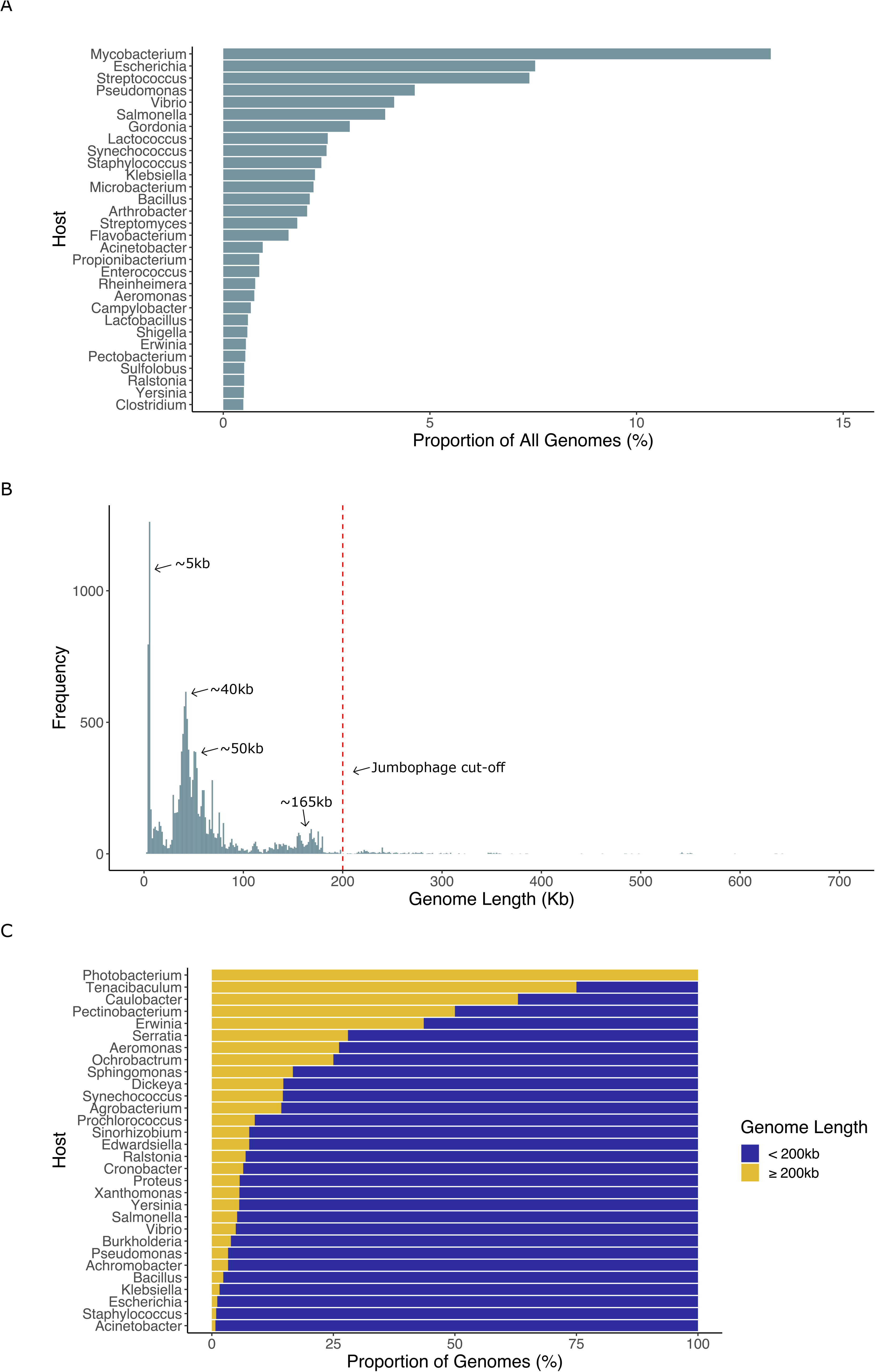
Overall properties of phages. A) Proportion of phages isolated on the top 30 most abundant hosts. B) Distribution of phage genome sizes. C) Proportion of “jumbo-phages” on top 30 hosts for which at least one “jumbo-phage” has been isolated.

**Figure 3.**
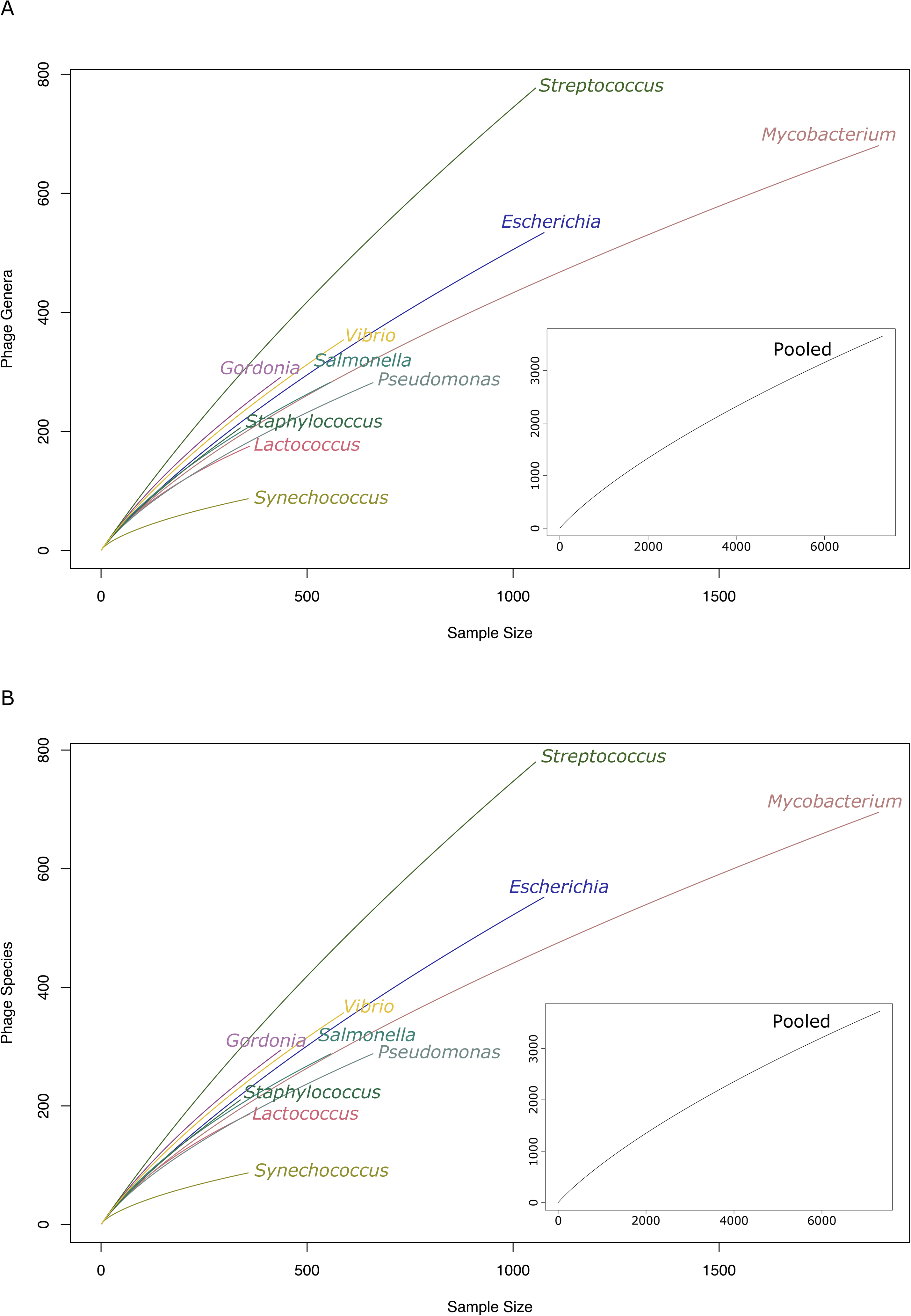
Genomic diversity of phages on the top ten most abundant hosts. A) Rarefaction curve of phage species. Species were defined as 95% identity over 95% of genome length. B) Rarefaction curve of phage genera. Genera were defined as 70% identity over 95% of genome length.

### Lytic and Temperate phages

To identify if the phage is lytic or temperate, we searched for genes that facilitate a temperate lifestyle (e.g., integrase and recombinase) that have been used in previous studies. This process is not perfect, as the presence of an identifiable gene linked to temperate phages does not mean it will access a lysogenic cycle. However, it does allow large scale comparative analyses, compared to the manual searching of literature of every phage compared to determine if it has been experimentally tested. Within the dataset, 4,258 (~30%) phages have the potential to access a lysogenic lifecycle. The frequency of putative temperate phages was highly variable depending on the host (Supplementary Figure 2). The number of putative temperate phages is also biased towards a small number of hosts with 1,217, 846 and 214 isolated on *Mycobacterium*, *Streptococcus* and Gordonia respectively. Collectively these three hosts account for ~54% of all putative temperate phage genomes sequenced to date (Supplementary Figure 2).

### Genomic Properties

Phage genomes ranged from 3.1 kb to 642.4 kb in size, with a clear distribution in the size of genomes with the most prominent peaks at 5-10 kb, 40 kb, 50 kb and ~165 kb (Figure 2b). The mean and median coding capacity was found to be 90.45% and 91.52%, respectively (Supplementary Figure 2). Of the 14,244 genomes, 5,731 (~40%) were found to have ≥ 90% of coding features on one strand and 3,293 (~23%) of these had coding features entirely on one strand (Supplementary Figure 2). The number of phages with genes encoding tRNAs was 4,590 (~32%). For those phages encoding tRNAs, the range was 1 to 62 with a median of 3. Whilst there is much literature on the presence of tRNAs in phages, it is still not clear entirely what role they provide to phages and why they absent in some phages and not others (46).

Phages with genomes greater than 200 kb are often referred to as “jumbo-phages” and are reported to be rarely isolated (47). 314 genomes (~2.2%) greater than 200 kb in length were identified, suggesting that they are rare. To further investigate if “jumbo-phages” are as rare as is thought, we looked at the distribution in the context of the previously identified host bias. “Jumbo-phages” have only been isolated on 31 of 234 identifiable bacterial hosts (Supplementary Table 1) and are far more commonly isolated on some hosts than others. Noticeably absent are any “jumbo-phages” that infect *Mycobacterium*, *Gordonia*, *Lactococcus*, *Arthrobacter*, and *Streptococcus*, with >4,000 phages having been sequenced from these bacterial hosts (Figure 2c). For host bacteria that have had far fewer phages isolated on them such as *Caluobacter*, *Sphingomonas*, *Erwinia*, *Areomonas*, *Dickeya* and *Ralstonia*, the frequency of “jumbo-phage” isolation is far higher (Figure 2c). Due to the small sampling depth of some of these hosts (e.g., *Photobacterium* and *Tenacibaclum*), it is not possible to determine whether the high proportion of genomes is merely a result of the low number of genomes sequenced. However, for other hosts such as *Aeromonas*, *Erwinia* and *Caulobacter* from which more than 20 phages have been isolated, ~26%, ~44% and ~63% are categorised as “jumbo” respectively. Therefore suggesting “jumbo-phages” are not always rare on particular hosts.

We further investigated the phylogeny of “jumbo-phages” using the translated sequence of the terL gene. The “jumbo-phages” are well distributed across the tree and do not form a single monophyletic clade, suggesting that they have arisen on multiple occasions, with multiple clades of phages having representatives of “jumbo-phages” within them. Not all “jumbo-phages” are equal, with “jumbo” cyanophages infecting the cyanobacteria *Synechococcus* and *Procholorococcus* only marginally larger than there non-jumbo cyanophages relatives. These “jumbo-phages” are also more closely related to their non-jumbo cyanophages relatives than other “jumbo-phages” (Figure 4). This is not limited to the cyanophages, with many other “jumbo-phages” more closely related to a non-jumbo phage. A similar pattern of grouping non-jumbo with “jumbo-phages” is observed when a reticulate approach is used to look at the relatedness of phage genomes using vConTACT2 (Supplementary Figure 3).

**Figure 4.**
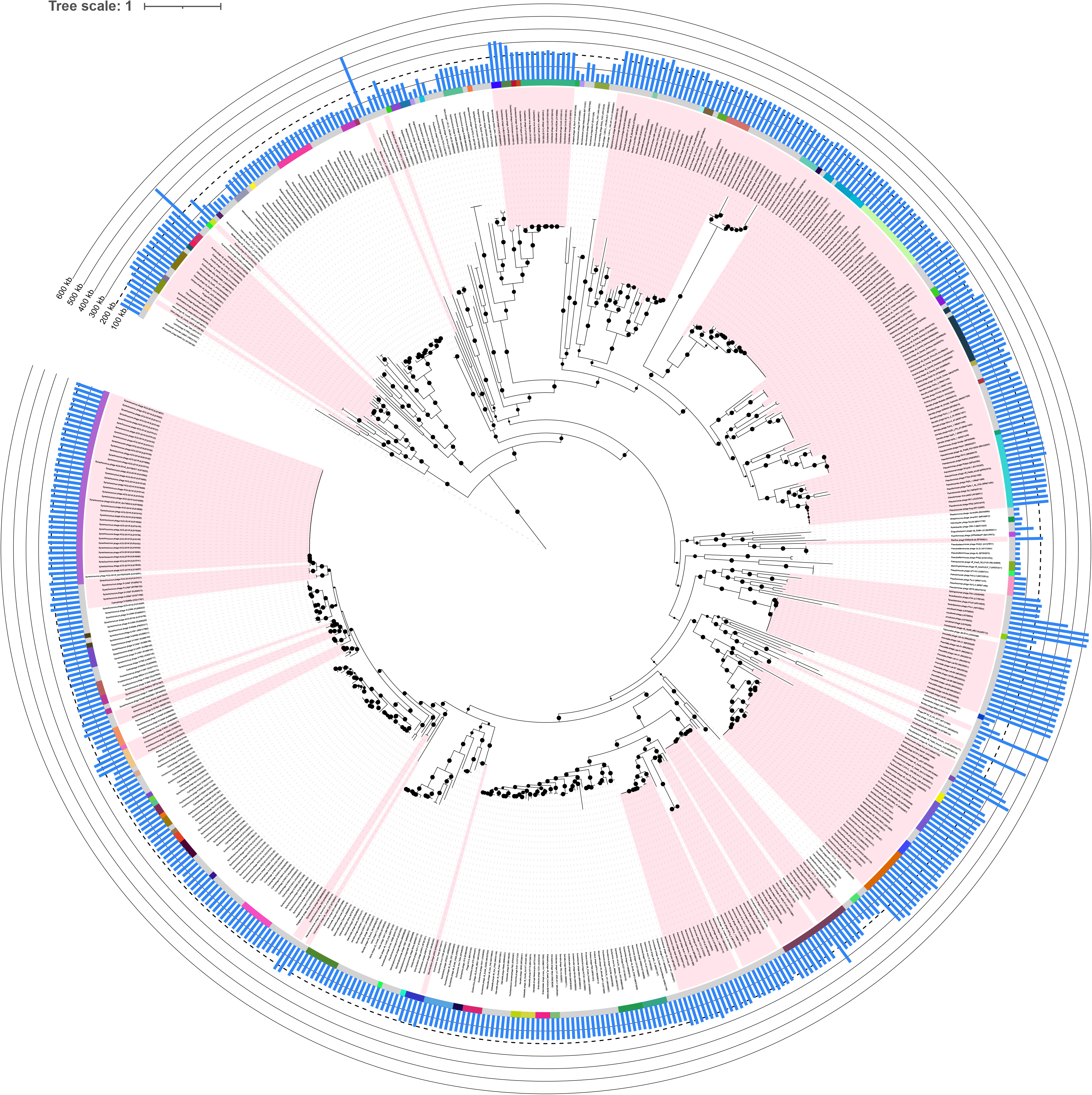
Phylogenetic tree of translated terL gene for 313 “jumbo-phages” and their closest relatives. The alignment was produced using MAFFT (36) and tree produced using IqTree using WAG model with 1000 bootstrap repeats (37). Pink shaded regions indicate “jumbo-phages”, coloured ring indicates viral genus, and blue bars indicate genome length. Bootstrap values indicated by black circles are shown with a minimum of 70%.

### Virulence Factors and Antimicrobial Resistance Genes

The presence of ARGs and virulence factors is major concern for phage therapy, as the use of phages carrying such genes may make the populations of bacteria they are intended to kill more virulent or resistant to antibiotics. We therefore used this database to integrate the frequency and diversity of phage-encoded virulence factors and ARGs. 235 genomes (~1.6%) were found to encode a virulence factor and 43 genomes (~0.3%) to encode an ARG. The most common virulence genes were the *stx*_2A_ (72 genomes) and *stx*_2B_ (71 genomes) genes that encode subtypes of the Shiga toxin (Supplementary Table 2). The most common ARGs were the *mef*(A) (14 genomes) and *msr*(D) genes which confer resistance to macrolide antibiotics (Supplementary Table 3) (48). Most genomes encoding a virulence factor were predicted to be from temperate phages (222/235), and were found to infect six bacterial genera, with the three most abundant hosts being *Streptococcus*, *Staphylococcus* and *Escherichia* respectively. The hosts for many genomes could not be determined (55/235). The virulence factor encoding genomes were widely distributed over 26 putative genera (Supplementary Figure 3). All genomes encoding an ARG were predicted to be temperate and were found to be isolated from eight bacterial genera, with the majority of phages linked to *Streptococcus* spp. (27/43).

## Discussion

Defining how many different complete phage genomes have been sequenced is not a simple question as it might appear. Based on accession numbers, there are 14,244 phage genomes, once RefSeq duplicates have been removed. Using unique names results in 12,127 phages, however using names alone does not give an accurate estimate of the number of different phages, as genomically different phages have the same name. The use of de-duplication at 100% identity suggests 13,830 unique phage genomes (January 2021) from cultured isolates. This assumes that the genome submissions are from isolates and not predictions of prophages from bacterial genomes. For the vast majority of phages, this appears to be case, although not easily discernible for all phage genomes.

The data reveals clear patterns in phage genomes and biases in the selection of phage genomes that are currently available, but not always discussed in the analysis of genomes. The first is the number of phage genomes is relatively small. Even for hosts where the highest number of phages have been isolated on, our estimates suggest 1000s of new phage species remain to isolated and sequenced. If we consider there are now more than 300,000 assembled representative bacterial genomes in GenBank, with many hundreds of thousands more for particular genera e.g., >300,000 *Salmonella* and *Escherichia* genomes alone (49). The representation of phage genomes to date is tiny compared to their bacterial hosts. Furthermore, the rate at which phage genomes are being sequenced is slowing down rather than increasing. Given the renewed interest in phages and increased accessibility of sequencing, the decrease in the rate over time was surprising.

The second point of note is the bias in phage genomes. With a clear bias in both the hosts phages are isolated on and for lytic phages over temperate phages. Thus, these phages are representative of these particular hosts, rather than phages in their entirety. Due to the enormous success of the SEA-PHAGES program, many phages have been isolated on *Mycobacterium* and *Gordonia* (50). This in turn results in ~1/3^rd^ of all temperate phage genomes being isolated on these two bacterial genera, whereas the remaining 2/3^rds^ are distributed across 142 different hosts.

The overrepresentation of phages infecting particular hosts can lead to truisms that may not be correct. For instance, “jumbo-phages”, those that have genomes >200 kb, are rarely isolated (47). Analysis of the complete dataset suggests ~2.2% of genomes fall into this category. However, this needs to be viewed in the context of the large bias in the hosts used for isolation, with ~75% of phages isolated on only ~16% of bacterial hosts that could be identified. When the number of “jumbo-phages” is expressed as a percentage of all phage genomes, their isolation is clearly rare. For some hosts, such as *Mycobacterium*, many hundreds of phages isolated on the same host strain have been sequenced without the isolation of a “jumbo-phage”, suggesting they are truly rare for this host (45). However, for other hosts such as *Procholorococcus*, *Synechococcus*, *Caulobacter*, and *Erwinia*, the isolation of “jumbo-phages” is not a rare event. While methodological adjustments of decreasing agar viscosity and large pore size filters may increase the number of phages isolated that have larger genome sizes (47), we suggest that using a wider variety of hosts may increase the number of “jumbo-phages” isolated. Phylogenetic analysis demonstrated many “jumbo-phages” are more closely related to non-jumbo phages than other “jumbo-phages”. Thus, as the number of phage genomes has increased an arbitrary descriptor or “jumbo” for phages with genomes over 200 kb in length has less meaning. Recent comparative analysis of 224 “jumbo-phages”, used proteome size and analysis of protein length to determine a cut-off of 180 kb to separate “jumbo-phages”, from other phages. From this using a clustering-based approach, three major clades of “jumbo-phages” were identified (51). In this study using *terL* as a phylogenetic marker to determine the phylogeny of 313 “jumbo-phages” and their closely related phages, suggests they have arisen on multiple occasions, as has been demonstrated previously (51). “Jumbo-phages” are clearly not monophyletic and what applies to one “jumbo-phage” does not hold true for many others (51). As the number and diversity of “jumbo-phages” increases, the use of the term seems to have less meaning.

With the increasing interest and use of phages for therapy, the isolation of phages that do not contain known virulence factors or ARGs is imperative. How frequently phages encode antibiotic resistance genes is a topic of much debate (52, 53). A previous study of 1,181 phage genomes found that they are rarely encoded by phages with only 13 candidate genes, of which four where experimentally tested and found to have no functional antibiotic activity (47). We estimate ~0.3% of phage genomes encode a putative ARG (none have been experimentally tested), a finding that is consistent with previous reports of low-level carriage in phage genomes (52) in a dataset that is ~10x larger using similarly stringent cut-offs. Critically, all of these ARGs were found in phages that are predicted to be temperate or have been engineered to carry ARGs as a marker for selection. With the frequency of carriage in temperate phages being ~1% overall. However, this data is still biased by the majority of temperate phages being isolated on only three bacterial genera. Notably no ARGs were detected on phages of *Mycobacterium*, which accounts for ~28 % of temperate phages. In comparison, ~2.6% (27/1055) of temperate phages of *Streptococcus* carry putative ARGs and 50% of phages from *Erysipelothrix* (1/2). Clearly a much deeper sampling of temperate phages from a broader range of hosts is required to get an accurate understanding of the role of phage in the carriage of ARGs. Based on the skewed data available to date, it seems unlikely there will be issues in the isolation of lytic phages for therapeutic use that contain known ARG within their genomes. However, we cannot determine whether these lytic phages cannot spread ARGs via transduction, or through carriage of as-yet uncharacterised ARGs.

Whilst there is much debate on the presence and importance of ARGs in phage genomes, the role of genes encoding virulence factors is well studied and the process of lysogenic conversion well known (7–10). However, how widespread known virulence genes are in phages is not widely reported. We estimate 1.6% of phages encode at least one putative virulence factor, with the frequency of carriage far higher in temperate phages (5.5%) than lytic phages (0.13%). Again, these overall percentages are skewed by host bias with no known virulence factors detected in *Mycobacterium* temperate phages (0/1217), in comparison 72% of temperate phages of *Shigella* (5/7) and 7% (61/846) of *Streptococcus* contain virulence factors. It is currently impossible to determine if the higher proportion of ARGs and virulence factors in phages of known pathogens is a feature of their biology, or a skew in the database towards phages of clinically relevant isolates.

Given the biases in the dataset, it is not clear if the general phage patterns we observe (e.g., jumbo-phages are rarely isolated, more temperate phages on particular hosts, and the carriage of ARGs and virulence genes) are linked to biology or chronic under sampling of phage genomes. We speculate that currently is most likely the latter, which distorts some generalisations about phages. It clear that jumbo-phages are not rare on some hosts and putative ARGs are far more abundant on temperate phages. However, far deeper sampling of phage diversity across different hosts is required at an increasing rate.

## Conclusions

We have provided a simple method to automate the download of curated set of complete genomes from cultured phage isolates, providing metadata in a format that can be used as a starting point for many common analyses. Analysis of the current data highlights what we know about phage genomes is skewed by the majority of phages having been isolated from a small number of bacterial genera. Furthermore, the rate at which phage genomes are being deposited is decreasing. Whilst understanding of genomic diversity is always influenced by the data available, this is particularly acute for phage genomes with so many phages isolated on smaller number of hosts. To obtain a greater understanding of phage diversity, larger numbers of phages, in particular temperate phages, isolated from a broader range of bacteria need to be sequenced.

## Authorship Confirmation Statement

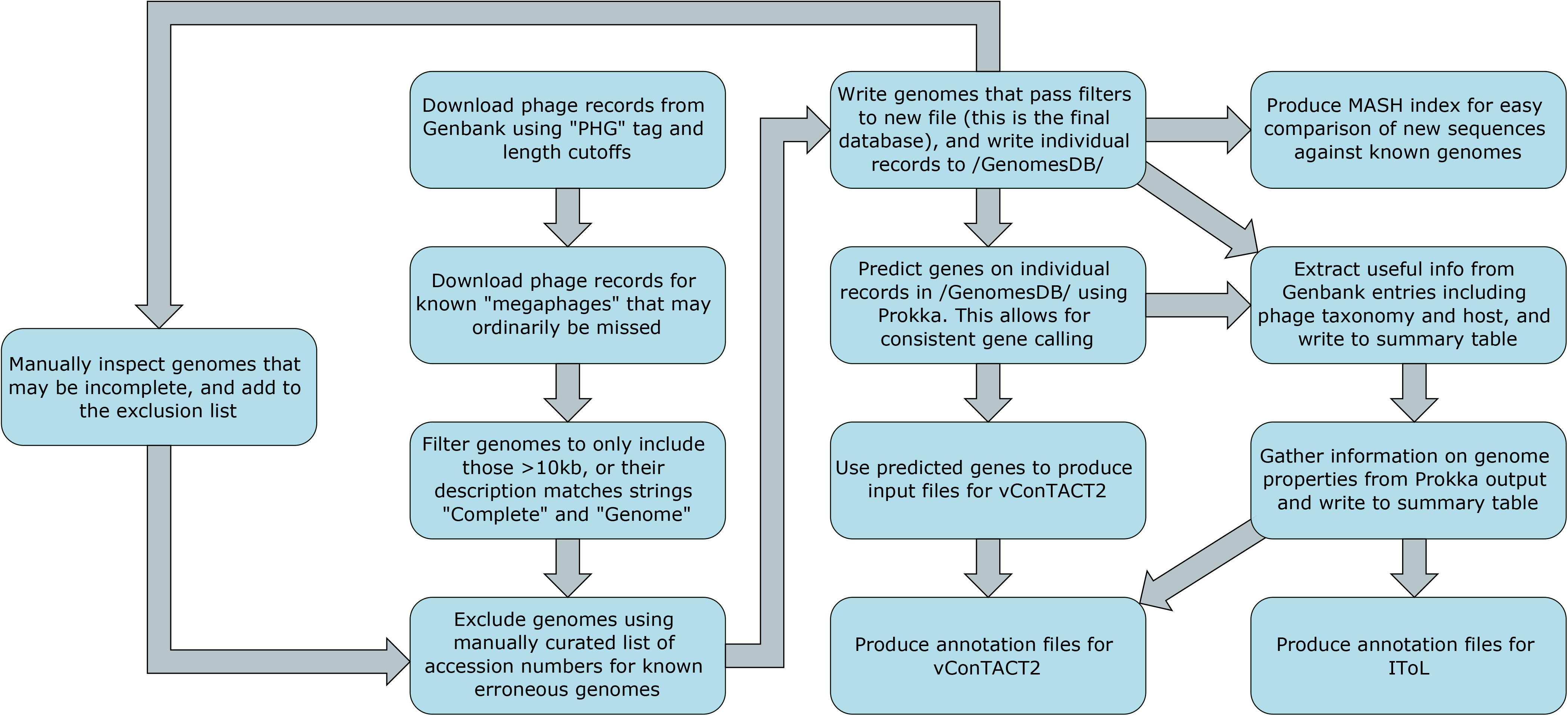

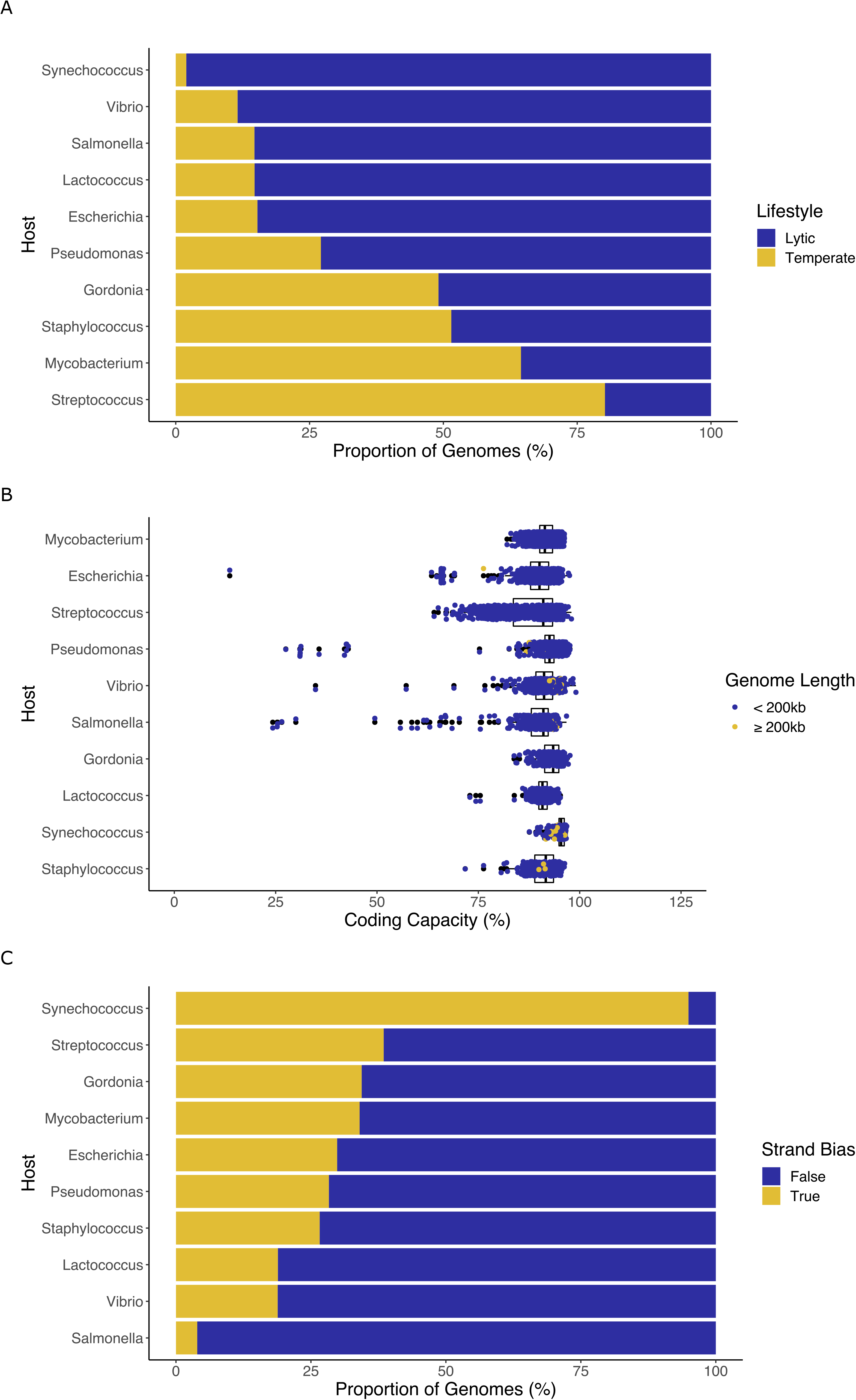

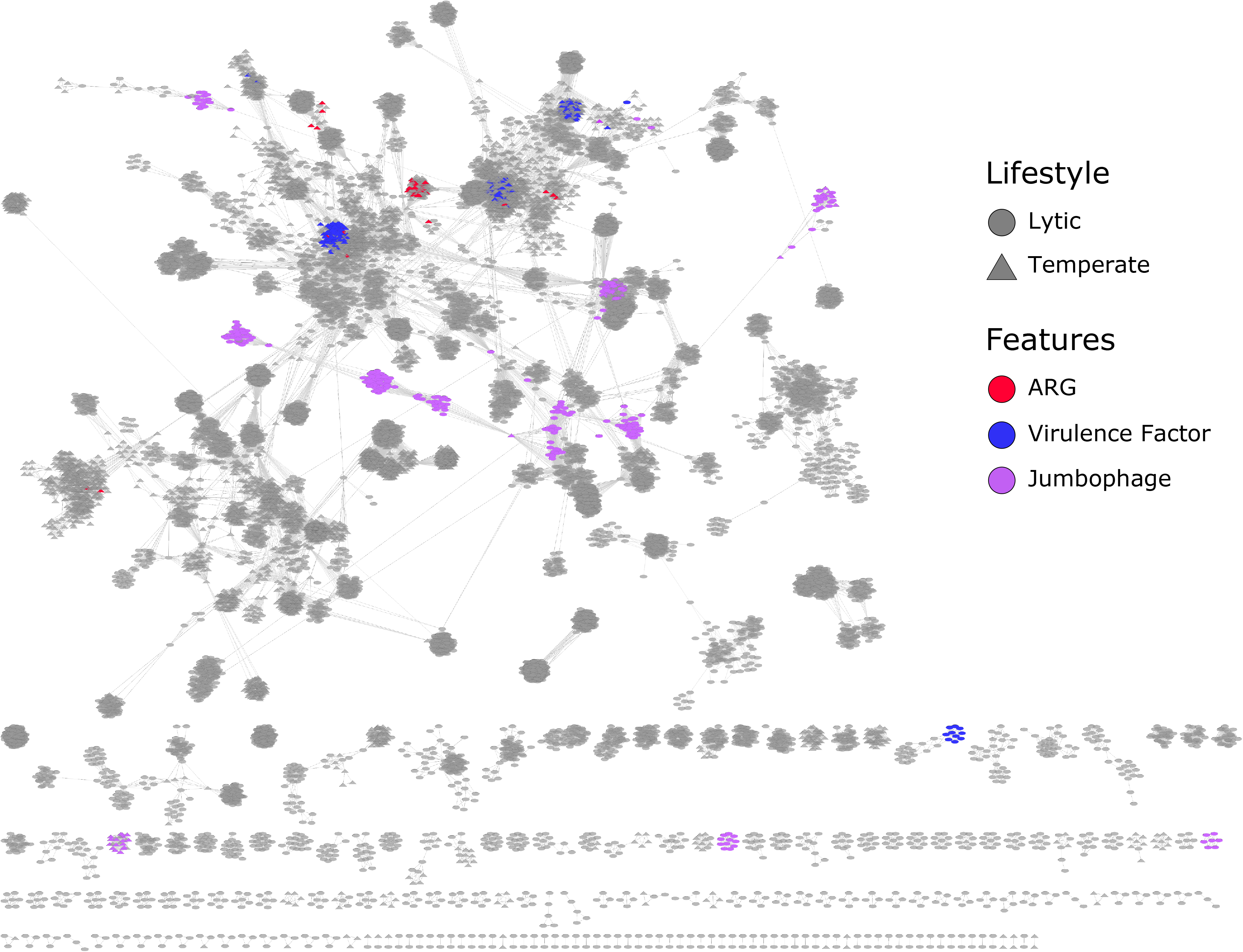

